# Conditional Denoising Diffusion Probabilistic Models with Attention for Subject-Specific Brain Network Synthesis

**DOI:** 10.1101/2025.01.06.631503

**Authors:** Meenu Ajith, Vince D. Calhoun

## Abstract

The development of diffusion models, such as Glide, DALLE 2, Imagen, and Stable Diffusion, marks a significant advancement in generative AI for image synthesis. In this paper, we introduce a novel framework for synthesizing intrinsic connectivity networks (ICNs) by utilizing the nonlinear capabilities of denoising diffusion probabilistic models (DDPMs). This approach builds upon and extends traditional linear methods, such as independent component analysis (ICA), which are commonly used in neuroimaging studies. A central contribution of our work is the integration of attention mechanisms into conditional DDPMs, enabling the generation of subject-specific 3D ICNs. Conditioning the resting-state fMRI (rs-fMRI) data on the corresponding ICNs enables the extraction of individualized brain connectivity patterns, effectively capturing within-subject and between-subject variability. Unlike prior models limited to 2D visualization, this framework generates 3D representations, providing a more comprehensive depiction of ICNs. The model’s performance is validated on an external dataset to prevent over-fitting and for overall generalizability. Furthermore, comparative evaluations also demonstrate that the proposed DDPM-based approach outperforms state-of-the-art generative models in producing more detailed and accurate ICNs, as validated through qualitative assessments.

## 1 INTRODUCTION

Diffusion models [1] have become the new benchmark in deep generative modeling [2], demonstrating exceptional performance across various applications. These generative models provide an alternative to existing generative adversarial networks (GANs) [3], variational autoencoders (VAEs) [4], and normalizing flow models [5]. For generative models to be utilized in real-world applications, they must meet several key criteria: high-quality sampling, diversity of samples, and fast, efficient sampling. While GANs can produce high-quality medical images rapidly, they often lack diversity and result in mode collapse [6]. In contrast, VAEs [7] and normalizing flows [8] excels at capturing the variation within data distributions but tend to produce lower-quality samples. Diffusion models address the limitations of VAEs and GANs by effectively capturing the variations in data distribution and generating high-quality samples. In recent years, denoising diffusion probabilistic models (DDPMs) [9, 10] have emerged as the leading algorithms for both unconditional and conditional image generation in the medical imaging domain. In the case of fMRI data, the application of high-fidelity fMRI image generation focused on 2D fMRI due to the computationally expensive nature of early diffusion models. A notable 2D medical data synthesis model [11] utilizes a DDPM framework integrated with a Swin vision transformer [12] to enhance the training set through augmentation. Later, another study proposed a 3D-DDPM for generating high-resolution MRIs of brain tumors [13]. Another alternative approach for the controllable generation of 3D MRI images is the use of latent diffusion models (LDMs) [14, 15]. This method produced 3D synthetic MRI images of the adult brain and utilized a cross-attention mechanism to integrate age, gender, and brain volume.

Blind source separation (BSS) [16, 17] is a fundamental problem in many image processing applications that aim at revealing latent structures within multivariate data through matrix decomposition techniques. Although the process is completely unaware of how the signals were mixed, it operates under certain assumptions about the source signals (e.g., the source distribution) or the mixing process (e.g., linear mixing). In terms of fMRI data, intrinsic connectivity networks (ICNs) are traditionally extracted using independent component analysis (ICA) [18, 19], a technique that decomposes the multivariate fMRI signals into spatially independent components that represent distinct brain networks. While ICA has been successfully applied in numerous studies, it remains challenging to manually select the relevant components and differentiate neurophysiological signals from noise. Other traditional BSS methods that have proven effective for neuroimaging data include sparse principal component analysis (sPCA) [20], sparse canonical correlation analysis (sCCA) [21], sparse partial least squares (sPLS)[22], and sparse two-dimensional canonical correlation analysis (s2DCCA) [23]. However, these traditional methods [24] are constrained by their linearity and inability to capture complex, non-linear relationships that may exist within the brain’s connectivity patterns.

Meanwhile, the success of deep learning across various neuroimaging applications has led researchers to explore its use in analyzing rs-fMRI data. Previous research applied restricted Boltzmann machines (RBMs) [25] to fMRI volumes to uncover intrinsic functional brain networks [26], resulting in a competitive performance compared to ICA. Another study [27] employed the RBM model on fMRI time courses to extract latent sources that distinguish internal neural activity within a single brain region and reveal functional interactions across multiple brain regions. This approach demonstrated higher correlations compared to ICA in identifying task-related components, though it required empirical parameter tuning. Additionally, researchers have proposed a framework that uses autoencoders and spatiotemporal sparsity constraints for blind source separation (BSS) from fMRI datasets [28]. While the sparse spatiotemporal BSS method shows promise with its computational simplicity and effectiveness, it can experience fluctuations during convergence and also faces model selection challenges. Therefore, despite these advancements, there remains a need for methods that are capable of generating high-quality, diverse, and efficient samples of ICNs from fMRI data.

Prior studies have demonstrated that DDPMs offer a promising alternative to GANs in the field of neuroimaging. DDPMs, known for their ability to iteratively refine noisy data into high-quality outputs, offer enhanced flexibility and precision in modeling complex connectivity patterns that traditional methods may overlook. This framework also addresses the challenge of noise and artifacts commonly present in fMRI data, improving the accuracy of ICN identification. By dynamically modeling individual differences in brain connectivity, DDPMs provide a more personalized and detailed representation of ICNs, facilitating deeper insights into brain function and variability. In this study, the proposed method establishes an initial framework by leveraging DDPMs for generating both 2D and 3D high-quality ICNs by harnessing the iterative refinement capabilities of DDPMs. Building upon this foundation, the framework advances by incorporating conditional DDPMs with cross-attention for generating subject-specific ICN generation from rs-fMRI data. By conditioning the individual’s rs-fMRI data, this approach ensures that the generated ICNs are highly tailored to the unique connectivity patterns of that subject. These DDPMs offer significant advantages for generating personalized ICNs by selectively focusing on relevant features within the rs-fMRI data. Thereby these models enhance the accuracy and detail of the generated ICNs while effectively managing noise and artifacts. Additionally, the model’s ability to adapt to individual differences and its efficient processing capabilities allow the model to capture both within-subject and between-subject differences of the ICNs. Consequently, this framework facilitates deeper insights into both the commonalities and individual variations in brain function, advancing our understanding of neuroscience and its applications in clinical settings.

The main contributions to this work are as follows:

- This work introduces a novel methodology for modeling brain connectivity, significantly contributing to the field by enabling the generation of clearer and more accurate representations of ICNs. Unlike traditional ICA-based methods, which are linear, DDPMs can capture non-linear relationships within brain connectivity patterns.
- This work applies DDPM to generate high-quality 2D and 3D intrinsic connectivity networks (ICNs) by capturing the underlying data distribution. This model is trained on a large dataset of ICNs obtained through the application of spatially constrained independent component analysis (ICA) to resting fMRI data.
- Generate subject-specific 3D ICNs by integrating attention mechanisms into conditional DDPMs. The model analyzes resting-state functional MRI (rs-fMRI) data conditioned on corresponding ICNs allowing it to learn unique brain connectivity patterns for individual subjects. During inference, the model accurately generates ICNs tailored to the specific rs-fMRI data provided, capturing both within-subject and between-subject variability.
- Comparative analyses demonstrate that the proposed DDPM model outperforms progressive GANs in generating ICNs. The DDPM model produces higher-quality images, as evidenced by qualitative assessments, highlighting its capability to better encapsulate the intricate details of brain connectivity networks. Additionally, the proposed approach can model and visualize both group-level and individual-specific brain connectivity patterns, offering new insights into the variability and complexity of brain function.

## 2 MATERIALS AND METHODS

### 2.1 Data Acquisition and Preprocessing of fMRI Data

The neuroimaging dataset for this analysis was obtained from the UK Biobank database [29], a large-scale dataset comprising older adults aged 53 to 87 years, with an average age of 69.75 ± 7.43 years. Resting-state fMRI (rs-fMRI) scans were performed using 3 Tesla (3T) Siemens Skyra scanners equipped with 32-channel head coils. The imaging protocol employed a gradient-echo echo planar imaging (GE-EPI) technique without iPAT, fat saturation, a flip angle (FA) of 52°, spatial resolution of 2.4×2.4×2.4 mm, field of view (FOV) of 88×88×64 matrix, repetition time (TR) of 0.735 s, echo time (TE) of 39 ms, and acquisition of 490 volumes. The scan duration was 6 minutes and 10 seconds, during which participants focused on a crosshair and stayed relaxed. Additionally, a multiband sequence with an acceleration factor of eight acquired eight slices simultaneously. Further to ensure high data quality, several preprocessing steps were applied to the UK Biobank data. Firstly, intra-modal motion correction was applied using the MCFLIRT tool [30]. Next, to enable comparison across participants, grand-mean intensity normalization was applied to scale the entire 4D dataset. High-pass temporal filtering addressed residual temporal drifts, and geometric distortions were corrected using FSL’s Topup tool [31]. Subsequently, EPI unwarping and gradient distortion correction (GDC) were conducted, and artifacts were removed using independent component analysis (ICA) combined with FMRIB’s ICA-based X-noiseifier [32]. The data were standardized to an MNI EPI template using FLIRT and SPM12, followed by the application of Gaussian smoothing with a full width at half maximum (FWHM) of 6 mm.

In this study, training is conducted using prior information input into the proposed DDPM model. This prior information consists of ICNs generated via spatially constrained ICA. We utilized NeuroMark [33], a fully automated spatially constrained ICA method, on the preprocessed rs-fMRI data. The NeuroMark_fMRI_1.0 template, which includes 53 ICNs derived from a 100-component blind ICA decomposition of two large healthy control datasets (the Human Connectome Project (HCP) and the Genomics Superstruct Project (GSP)), served as a template in an adaptive ICA approach. This approach enabled the estimation of subject-specific functional networks and their time courses (TCs). Based on their anatomical and functional characteristics the network templates were classified into seven functional domains, such as subcortical (SC: 5 ICNs), auditory (AUD: 2 ICNs), sensorimotor (SM: 9 ICNs), visual (VIS: 9 ICNs), cognitive control (CC: 17 ICNs), default mode (DM: 7 ICNs), and cerebellar (CB: 4 ICNs) domains.

### 2.2 Methods

#### 2.2.3 Denoising Diffusion Probabilistic Models

DDPMs form the foundation of current diffusion models in generative AI. They utilize a diffusion-derived approach to learn the underlying distribution of the data. These models have gained popularity due to their ability to generate high-fidelity samples and their stability during training. They generate these complex data distributions by iteratively refining random noise. The central idea behind DDPMs is that they are based on a diffusion process that systematically adds noise to data in a forward process and learns to reverse this process during generation. DDPMs mainly operate in two distinct modes. In the unconditional mode, images are generated from noise without any external input. In the case of conditional DDPMs, additional information is incorporated to guide the image generation process. This additional information involves class labels, textual descriptions, or other forms of structured data, allowing for the generation of data adhering to specific conditions.

The forward process in DDPMs is a Markov chain where Gaussian noise is added to the data sample **x**_0_ over a fixed number of timesteps *T*. This process continues till the data becomes indistinguishable and becomes pure Gaussian noise by time *T*. The forward process is also parameterized by a noise variance scheduler *β* that controls the noise variance at each timestep. Mathematically, the forward process is defined by a noising function, which is a conditional distribution *q* (**x**_*t*_ |**x**_(*t* −1)_). This function determines how noise is added at each step based on the previous time step’s image.

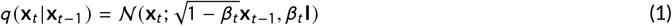

where 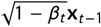 represents the mean *µ*_*t*_, and *β*_*t*_ **I** denotes the variance **Σ**_*t*_. As *t* approaches *T*, **x**_*t*_ increasingly resembles a sample from a Gaussian distribution 𝒩 (0, **I**). Moreover, the forward process can also be expressed in a closed form using the reparameterization trick as follows:

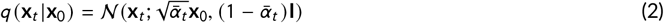

where we define 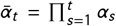 and *α*_*t*_ = 1 − *β*_*t*_. This allows for efficient sampling from any intermediate step *t* given the original data **x**_0_. The variance parameter *β*_*t*_ can be constant or vary over *T* timesteps. The original DDPM used a linear schedule, increasing *β* from 10^−4^ to 0.02 at the end.

Following this, during the reverse process, an image-generating model is used to predict the noise added to an image at a specific timestamp. Here the original data **x**_0_ is recovered from the noisy observation **x**_*T*_ by sequentially denoising it over the same *T* steps. The reverse process is modeled using the conditional distribution function *p* (**x**_*t* −1_) |**x**_*t*_). Since directly modeling this function is infeasible due to the numerous possible values of **x**_*t* −1_ for a given **x**_*t*_, neural networks are employed for this estimation. The function is modified to *p*_*θ*_ (**x**_*t* −1_) |**x**_*t*_) where *θ* represents the network parameters.

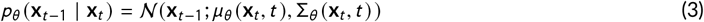

The training of this network is done by optimizing the variational bound on the negative log-likelihood of the data.

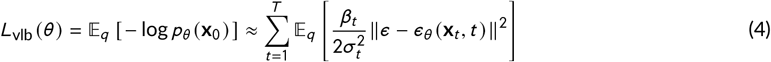

The simplified training objective for DDPMs is expressed as follows:

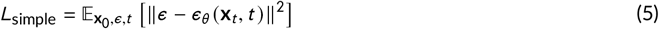

where *ϵ* is the Gaussian noise added at each step, and *ϵ*_*θ*_ is the neural network’s estimate of this noise.

In conditional DDPMs, the reverse process is extended to incorporate conditioning information **y**, which could represent any additional data such as labels or other modalities. The conditioning allows the model to generate samples that align with the given context, thereby enabling a controlled and targeted generation of outputs. The conditional reverse process is thus defined as:

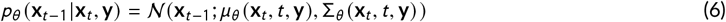

where the mean and variance also depend on the conditioning variable **y**. The objective function for training the conditional model is similarly modified to include the conditional variable:

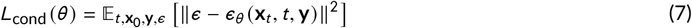

In medical imaging, conditional DDPMs have several key advantages. These include high-quality image generation from noisy or limited data, which aids in diagnosis and treatment. They provide personalized applications by conditioning the generative model on labels or textual descriptions. These models are robust to noise and can therefore be used to enhance low-resolution and corrupted medical images. They also support data augmentation and can be applied in multimodal applications, such as combining MRI with anatomical labels or synthesizing data across different modalities. The main strength of conditional DDPMs lies in their ability to combine the robustness of diffusion-based generative models with the flexibility of conditional generation, resulting in high-quality, diverse, and controllable outputs.

#### 2.2.2 ICN generation framework

The proposed DDPM model includes two distinct frameworks: the ImageGen Framework for 2D and 3D ICN generation, and the CondAttn-DDPM Framework, which leverages conditional DDPMs with attention mechanisms to generate subject-specific ICNs.

The ImageGen Framework modifies DDPMs to effectively generate 2D and 3D neuroimaging data. For 2D neuroimaging, the framework processes individual slices of images, leveraging a U-Net-like architecture that includes encoder-decoder structures with skip connections to capture detailed features and context. For 3D data, the framework employs 3D convolutional layers to handle volumetric data, enabling the model to understand spatial relationships in three dimensions. The diffusion process involves progressively adding noise to the neuroimaging data and training the model to reverse this process, reconstructing high-quality images from noisy inputs. The model uses a loss function to ensure that generated images closely match real neuroimaging data, making it versatile for both 2D and 3D applications and suitable for generating synthetic data for diagnostic and visualization purposes. The ImageGen Framework employs standard DDPMs for general random image generation, producing images without incorporating any specific context or conditions. They learn to generate a wide variety of images from a broad dataset without focusing on any subject’s unique characteristics.

In contrast, the CondAttn-DDPM Framework is tailored for subject-specific generation, where the model incorporates additional information or conditions to generate images that are tailored to individual subjects. This conditional approach allows the model to produce neuroimaging data that reflects specific attributes of subjects, resulting in more personalized and contextually relevant outputs. Specifically, this framework conditions the ICNs against the corresponding rs-fMRI data and uses an attention mechanism to process both data types during training. During inference, the model is provided only with the rs-fMRI data, and it generates the corresponding ICNs for the subjects based on the learned relationships. In this framework, cross-attention plays a vital role in effectively integrating and utilizing both rs-fMRI data and ICNs. This mechanism allows the model to focus on the relevant features and relationships between these two types of data, enhancing its ability to capture complex dependencies and interactions. The cross-attention mechanism improves representation learning, filters out noise, and ensures that the generated ICNs are both accurate and contextually relevant. Hence, by leveraging subject-specific information and attention mechanisms, this model produces more accurate and meaningful neuroimaging data, making it highly effective for personalized diagnostics and targeted research.

In terms of architecture, while the forward process is consistent across both frameworks, the reverse process differs between them. In the ImageGen Framework, the U-Net architecture features three down-sampling stages, each incorporating a 2D or 3D convolutional layer followed by max-pooling to reduce the spatial dimensions of the feature maps. This is followed by a central bottleneck with several convolutional layers that further decrease the number of feature maps. After the bottleneck, the architecture includes three upsampling stages, each involving a transposed convolutional layer and concatenation with the corresponding downsampled feature maps. During image generation, the model begins with a noisy image and iteratively refines it by learning to denoise step-by-step. It takes the image at time *t* and the timestep *t* as inputs and predicts the noise present in the image. By progressively removing the noise and reconstructing the original image, the model generates high-quality, detailed images from initial random noise.

The CondAttn-DDPM architecture is shown in Fig. 1 and it utilizes a sophisticated design to integrate subject-specific information through cross-attention mechanisms, enhancing the generation of neuroimaging data. The model begins by incorporating sinusoidal embeddings to encode time steps, which are then projected into a higher-dimensional space using time embedding layers. This process encodes temporal information, enabling the model to differentiate between various stages of the denoising process. The first half of the network consists of several down-sampling blocks. Each block applies 3D convolutions and normalization, followed by max-pooling operations. This helps to reduce the spatial dimensions and extracts features at different levels. These sequential blocks progressively refine the input features, capturing increasingly abstract representations. The bottleneck phase incorporates a multi-head attention mechanism to integrate and align features from the mid-level representations with the projected context data (rs-fMRI data). This mechanism enables the model to focus on relevant features from the context while processing the main features. This approach allows the model to effectively use subject-specific data during inference. In the second half of the network, up-sampling layers progressively reconstruct the spatial dimensions of the output, combining features from previous layers to generate high-quality, subject-specific neuroimaging data. The final output is passed through a 3D convolutional layer, which further refines the generated image.

**FIGURE 1.**
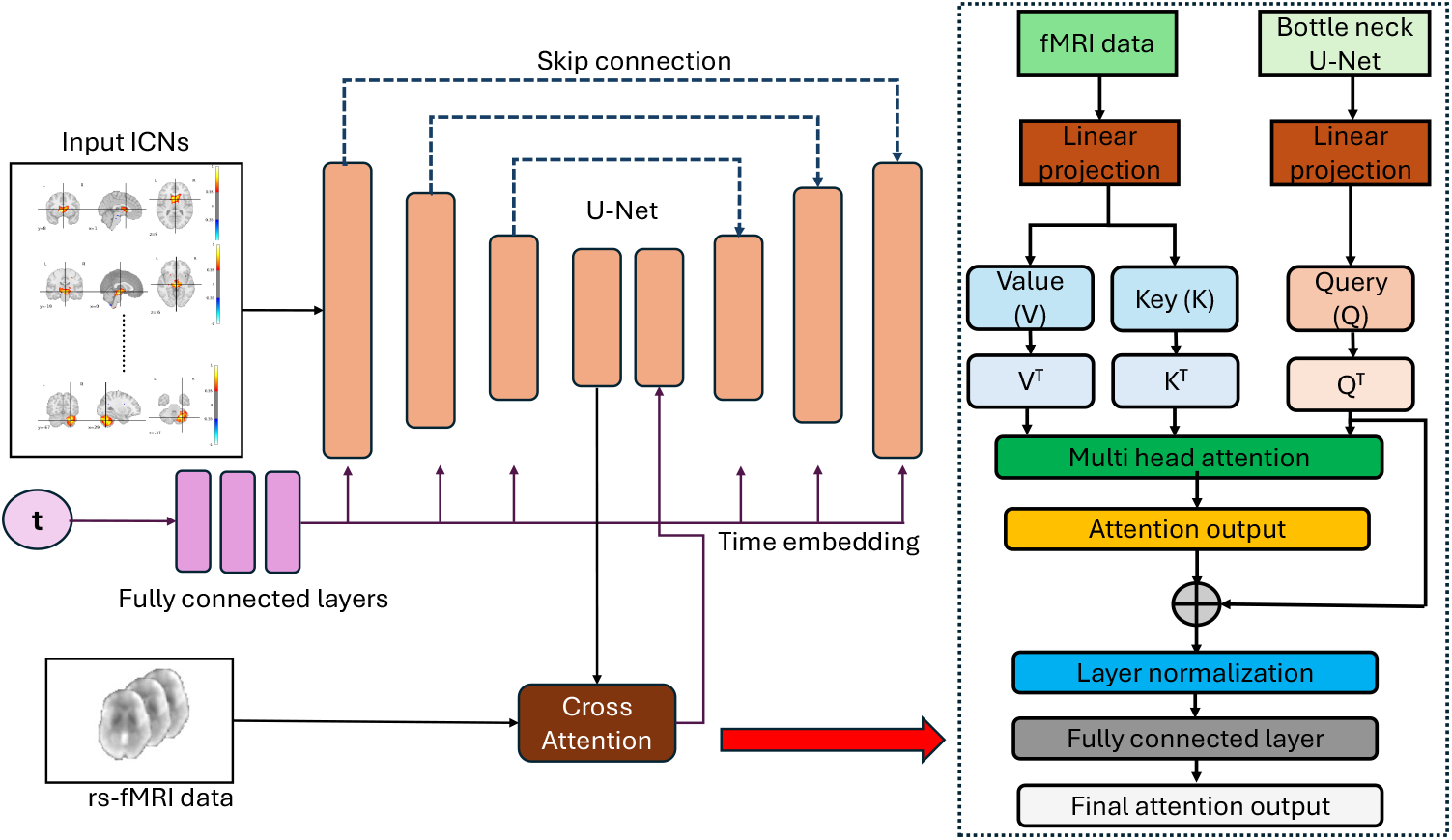
The architecture of the proposed CondAttn-DDPM model.

#### 2.2.3 Cross-attention mechanism and conditionality

In the CondAttn-DDPM architecture, cross-attention helps integrate information from different sources, such as the ICNs and the rs-fMRI data. The process involves projecting the input features and context features into a common embedding space and then applying attention to align and integrate them. Let *Q* be the query (ICNs), *K* and *V* be the key and value matrices (rs-fMRI data), and *W*_*Q*_, *W*_*K*_, *W*_*V*_ be the linear projection matrices.

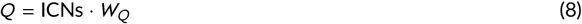

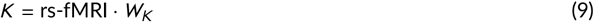

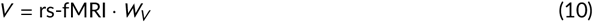

Here conditionality helps to ensure that the generated neuroimaging data reflects the specific attributes of the subject by incorporating additional information into the generation process. In this architecture, the rs-fMRI data is used to condition the generation of ICNs. During the conditioning process, the rs-fMRI data is projected into the same embedding space as the ICNs. This transformation allows the model to match the feature from both data sources. It helps to map the high-dimensional rs-fMRI data into a lower-dimensional embedding space. This makes it compatible with the ICNs during the cross-attention mechanism. During the bottleneck phase, the model uses the cross-attention mechanism to integrate the rs-fMRI embeddings with the ICNs. This is done by computing the dot product between *Q* and *K*. The result is then passed through a softmax function to obtain a weight distribution that indicates the importance of each key for the corresponding query. These weights are then used to compute a weighted sum of the values, which produces the attention output. The query is then added to the attention output, and layer normalization is applied to it. It helps to improve convergence during training by normalizing the output across the features. This is followed by a fully connected layer to obtain the final attention output. This layer helps refine the features learned through attention so that the output can be further processed in subsequent layers of the model.

The model is trained using a loss function that measures the difference between the predicted noise and the true noise added during the forward diffusion process. This loss function ensures that the model learns to accurately reverse the noise addition, leading to high-quality image generation. Here the model takes as input a noisy image along with a time step embedding and conditioning data. This conditioning ensures that the generated ICNs capture the specific traits of the subject as represented in the rs-fMRI data.

The conditioning data is used to guide the generation process, ensuring that the output is tailored to subject specific characteristics.

## 3 EXPERIMENTAL SETUP AND RESULTS

The proposed DDPM model, consisting of two frameworks, the ImageGen Framework and the CondAttn-DDPM Framework, is trained on rs-fMRI data from 10,000 subjects in the UK Biobank database. Preprocessing steps such as normalization and background removal are applied before training to enhance the quality and consistency of the input data. Normalization helps standardize the data by scaling it to a consistent range, reducing variability that could arise from differences in the intensity of the input images. Meanwhile, background removal eliminates irrelevant parts of the image, such as noise and artifacts, allowing the model to focus on the brain’s functional networks. For each subject, 53 ICNs are generated using ICA, with each ICN representing different functional domains of the brain. By leveraging this large and diverse dataset, the model successfully captures the variability across different networks. It learns to generate realistic images that reflect the unique patterns of brain activity representing each ICN.

During training, the data is processed in batches of 32 images to optimize memory usage, and the model is trained over 20 epochs. The DDPM model uses 1000 discrete time steps in which noise is gradually added to the data. The beta schedule ranges from 10^−4^ to 0.02, controlling the variance of the noise at each step. During each epoch, for each batch of images, the model introduces random noise to the images (forward diffusion process) and then attempts to reverse this noise (backward diffusion process) to recover the original image. This is achieved using the Mean Squared Error (MSE) loss function that compares the model’s predicted noise with the actual noise added to the images. The optimization process uses the Adam optimizer with a learning rate of 0.001 to adjust the model’s weights to minimize the loss. Hence the overall goal of the model is to predict the noise that was added to the images during each time step. At the end of each epoch, the average loss is recorded, and the model is saved if it achieves a new lowest loss. Once training is complete, the model’s performance is evaluated on the Human Connectome Project (HCP) [34] dataset, which serves as an independent test set. The best-performing model, identified during training, is then used for validation, ensuring that the model can effectively generalize across different datasets. The validation process is similar to training but does not involve updating the model weights. The goal here is to measure how well the model can generate new images based on the reverse diffusion processes it learned during training. For subject-specific generation, the training process utilizes both rs-fMRI data and their corresponding ICNs. In contrast, during validation, only the rs-fMRI data is provided as input, allowing the model to generate the corresponding ICNs.

### 3.1 2D ICN generation: A comparative analysis with ICA and Progressive GANs

The performance of 2D ICN generation was assessed using different models such as spatially constrained ICA, progressive GANs, and 2D-DDPMs. This comparison revealed notable differences among the models, both in terms of visual output and computational performance, as shown in Fig. 2. A visual analysis of the seven different ICNs generated by each model has provided clearer insights into their efficiency and output quality. Although ICA is not a generative model like GANs or DDPMs, it performed well in capturing certain structural elements of ICNs. It was particularly effective at identifying and capturing linear relationships in the data, resulting in images that accurately reflected simple and uniform patterns present in the ICNs. However, ICA struggled with more complex, non-linear relationships found in intricate brain networks. This limitation arises from ICA’s design, which focuses on independent components and linear separation. Meanwhile, progressive GANs, although known for generating high-resolution images, performed poorly when tasked with generating 2D ICNs. The visual representation of the generated ICNs highlighted a significant number of artifacts and random noise. It also suffered from mode collapse, a common issue in GANs where the model fails to capture the diversity of the training data, instead producing limited variations in the output. This was observed through the repetitive patterns seen in several of the generated ICNs, which failed to capture the complexity and diversity of the ICNs.

**FIGURE 2.**
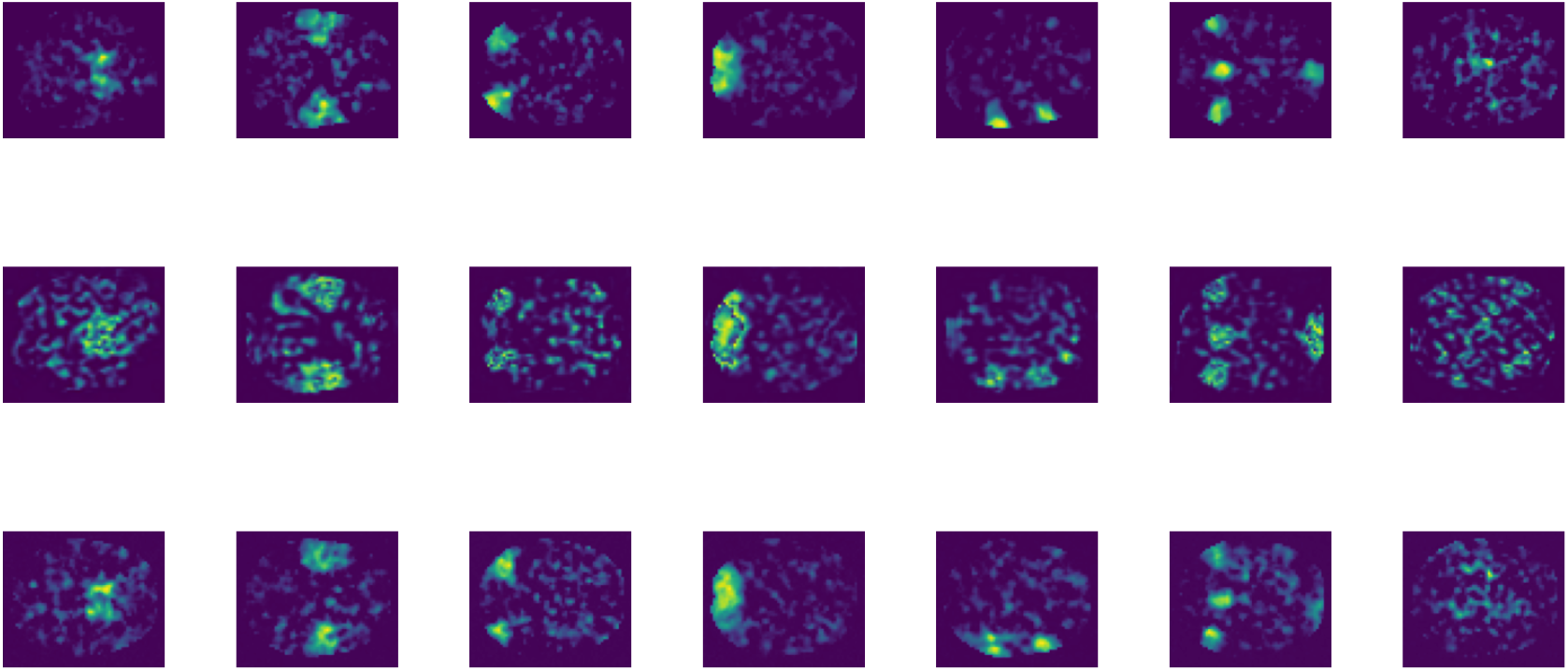
2D-ICN generation showcasing different methods: the first row presents ICNs generated using ICA, the second row displays ICNs generated by a Progressive GAN model, and the third row features ICNs produced using the 2D-DDPM Model.

Finally, the ICNs generated by the 2D-DDPM model were notably superior, effectively capturing both linear and non-linear dependencies within the data. The model’s iterative, step-by-step denoising process resulted in highly detailed and distinct ICNs. Unlike progressive GANs, the outputs displayed no visible artifacts and noise, which is critical for tasks requiring high precision. Moreover, in contrast to Progressive GANs, which experienced mode collapse, DDPMs consistently produced diverse and unique ICNs across multiple iterations, showcasing their ability to avoid common generative model challenges while maintaining output variability. Furthermore, an analysis of the computational efficiency and training times of the three models revealed significant differences that affect their practical applicability for 2D ICN generation. DDPMs exhibited an efficient training process, requiring approximately 40 minutes to complete 20 epochs. In contrast, progressive GANs required a much longer training period, taking about 2 hours and 47 minutes for the same number of epochs. This computational burden can be attributed to the complexity of their architecture and the progressive nature of their training approach. Conversely, ICA was less computationally demanding than the other two methods, utilizing simpler calculations for fMRI decomposition, resulting in faster training times. However, this speed comes at the cost of output quality and complexity, as ICA struggles to model the non-linear relationships present in ICNs. Overall, DDPMs provide a good balance of quality and efficiency compared to the rest of the models.

### 3.2 3D ICN generation and validation on an external dataset

The ICN images generated using 3D-DDPMs are illustrated in Fig. 3. A single-subject rs-fMRI data generates 53 ICNs, with each of them belonging to seven different functional domains such as subcortical, auditory, sensorimotor, visual, cognitive control, default mode, and cerebellar domains. 3D-DDPMs are advanced generative models used to iteratively learn complex distributions by gradually denoising data from a noise process.

**FIGURE 3.**
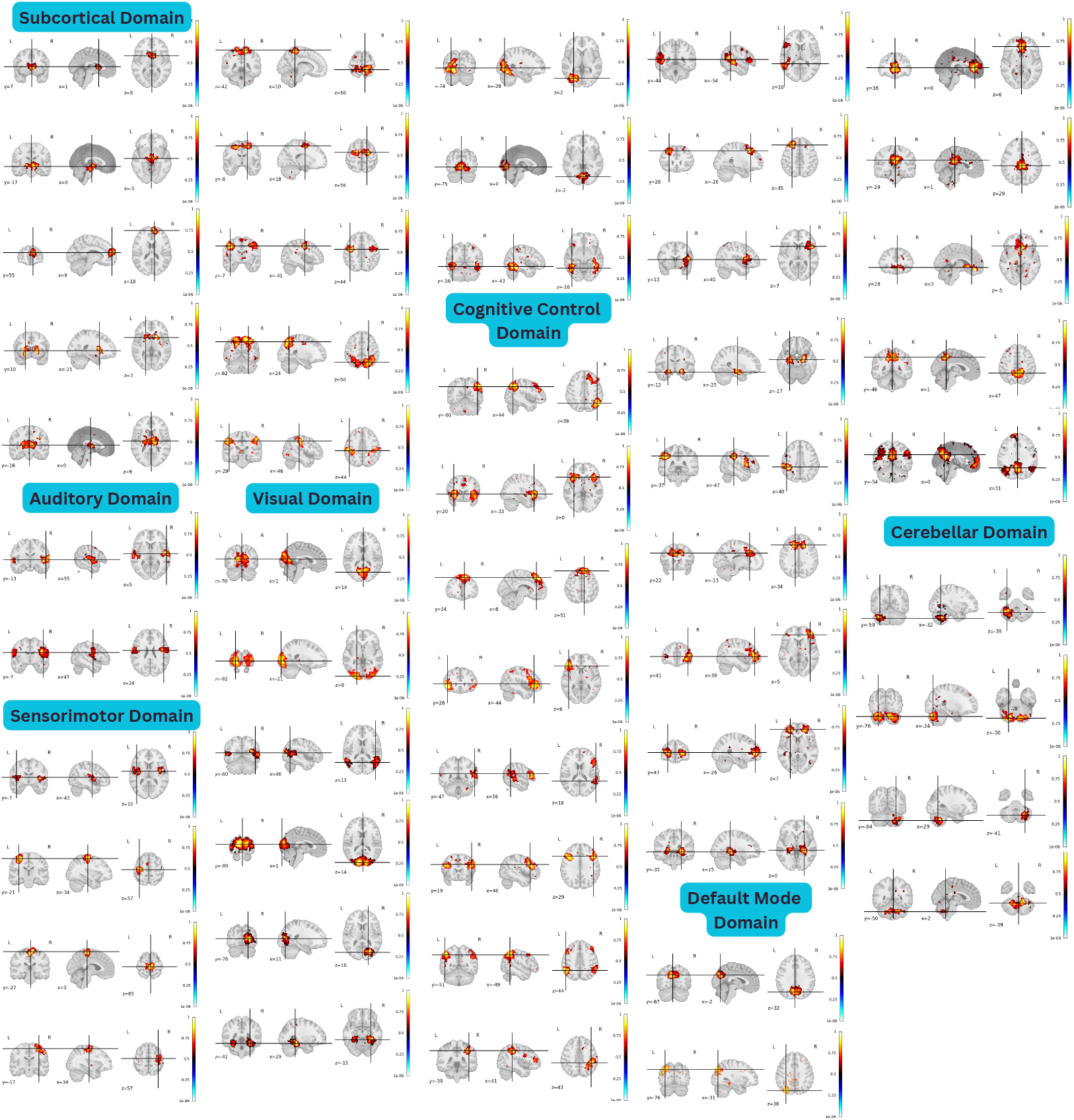
3D-ICN generation from single-subject rs-fMRI data using 3D-DDPMs. The model produces 53 ICNs, grouped into 7 distinct functional domains.

In this case, the 3D-DDPMs generate spatially meaningful ICNs from rs-fMRI data by capturing the spatio-temporal correlations in brain networks. These ICNs were validated on the HCP dataset by comparing them with ICNs derived using ICA. Here we utilized 500 generated images from each of the models for this comparative analysis. Quantitative metrics yielded a mean MS-SSIM of 0.72, MSE of 0.001, and PSNR of 27 across the generated ICNs, as shown in Table. 1. While these values indicate reasonable similarity to ICA-generated networks, it is important to note that very high MS-SSIM or PSNR, or very low MSE, is not indicative of better performance during this comparison. ICA is a linear decomposition method that assumes statistical independence, whereas the 3D-DDPM is a deep generative model designed to capture complex, nonlinear relationships that linear methods like ICA may miss. The observed metrics reflect a moderate degree of overlap with ICA-derived ICNs but also emphasize the added benefit of 3D-DDPMs in modeling additional subject variability and possible nonlinearities of rs-fMRI data. These differences highlight the model’s ability to capture richer and more detailed connectivity patterns that better reflect the inherent complexity of brain networks.

**TABLE 1.** Validation of the 3D-DDPM model on external dataset.

| ICNs | HCP dataset validation |  |  | ICNs | HCP dataset validation |  |  |
| --- | --- | --- | --- | --- | --- | --- | --- |
|  | MS-SSIM | PSNR | MSE |  | MS-SSIM | PSNR | MSE |
| ICN 1 | 0.75 | 29 | 0.001 | ICN 28 | 0.66 | 26 | 0.002 |
| ICN 2 | 0.72 | 27 | 0.002 | ICN 29 | 0.74 | 27 | 0.002 |
| ICN 3 | 0.75 | 27 | 0.002 | ICN 30 | 0.72 | 26 | 0.002 |
| ICN 4 | 0.71 | 28 | 0.001 | ICN 31 | 0.67 | 26 | 0.002 |
| ICN 5 | 0.68 | 27 | 0.002 | ICN 32 | 0.75 | 26 | 0.002 |
| ICN 6 | 0.72 | 27 | 0.002 | ICN 33 | 0.68 | 27 | 0.002 |
| ICN 7 | 0.71 | 26 | 0.001 | ICN 34 | 0.71 | 26 | 0.002 |
| ICN 8 | 0.78 | 28 | 0.001 | ICN 35 | 0.72 | 26 | 0.002 |
| ICN 9 | 0.72 | 27 | 0.001 | ICN 36 | 0.69 | 26 | 0.003 |
| ICN 10 | 0.73 | 28 | 0.002 | ICN 37 | 0.72 | 28 | 0.001 |
| ICN 11 | 0.67 | 27 | 0.002 | ICN 38 | 0.68 | 26 | 0.002 |
| ICN 12 | 0.73 | 27 | 0.002 | ICN 39 | 0.80 | 29 | 0.001 |
| ICN 13 | 0.70 | 27 | 0.002 | ICN 40 | 0.70 | 27 | 0.002 |
| ICN 14 | 0.74 | 27 | 0.002 | ICN 41 | 0.68 | 26 | 0.002 |
| ICN 15 | 0.70 | 26 | 0.002 | ICN 42 | 0.73 | 28 | 0.001 |
| ICN 16 | 0.75 | 27 | 0.001 | ICN 43 | 0.77 | 29 | 0.001 |
| ICN 17 | 0.70 | 26 | 0.002 | ICN 44 | 0.72 | 27 | 0.002 |
| ICN 18 | 0.70 | 26 | 0.001 | ICN 45 | 0.70 | 27 | 0.002 |
| ICN 19 | 0.73 | 27 | 0.001 | ICN 46 | 0.72 | 26 | 0.002 |
| ICN 20 | 0.71 | 27 | 0.001 | ICN 47 | 0.76 | 29 | 0.001 |
| ICN 21 | 0.75 | 27 | 0.001 | ICN 48 | 0.68 | 26 | 0.002 |
| ICN 22 | 0.75 | 27 | 0.002 | ICN 49 | 0.69 | 28 | 0.002 |
| ICN 23 | 0.70 | 27 | 0.001 | ICN 50 | 0.72 | 29 | 0.001 |
| ICN 24 | 0.70 | 27 | 0.002 | ICN 51 | 0.73 | 26 | 0.002 |
| ICN 25 | 0.70 | 27 | 0.002 | ICN 52 | 0.77 | 28 | 0.001 |
| ICN 26 | 0.80 | 27 | 0.001 | ICN 53 | 0.67 | 26 | 0.002 |
| ICN 27 | 0.70 | 27 | 0.002 | Average | 0.72 | 28 | 0.001 |

### 3.3 Visualizing diffusion and probability flow lines

A comprehensive visualization of the reverse diffusion process focusing on the transformation of pixel intensity distributions over time is shown in Fig. 4. An overlapping plot is analyzed to capture the denoising trajectory, demonstrating the progression from a Gaussian prior to the generated data distributions. The y-axis in this plot represents the timesteps, ranging from *T*, which in this case is 1000, down to 0. This time-reversed progression captures the core dynamics of the denoising diffusion process.

**FIGURE 4.**
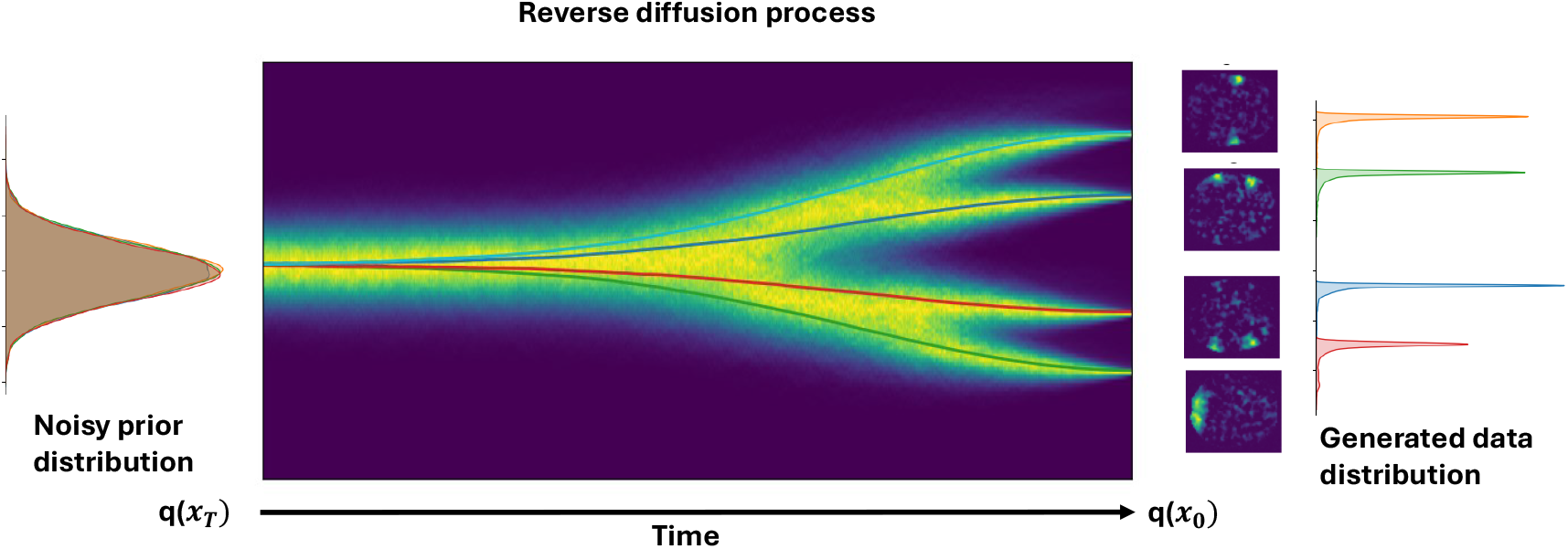
Visualization of the reverse diffusion process modeled as a stochastic differential equation (SDE). The process begins with a noisy prior distribution (left), typically a Gaussian, and iteratively denoises to produce a generated data distribution (right) that approximates the true data distribution.

Formerly, for each time point in the reverse process, the mean pixel intensity across the different images is computed. These images are generated iteratively during the reverse process, starting from random Gaussian noise and progressively converging to coherent data structures. The mean intensity for each time point is then plotted, and this represents how pixel intensities change as the denoising progresses. Hence this visualization highlights the temporal dynamics of intensity variations by offering insights into the convergence of the generated images. Additionally, histograms are also generated at each time point, demonstrating the distribution of pixel intensities over time. These histograms are arranged in a matrix format, and they provide a detailed visual representation of how pixel intensities evolve for each image throughout the reverse process.

The primary inference from these visualizations is that, during the reverse process, by sampling from the Gaussian distribution, the model generates new images that align with the underlying statistical properties of the original data. While these generated images are not exact replicas of the original data, they reflect the diversity and variability inherent in the learned data distribution. Here the left side of the plot corresponds to the initial Gaussian prior distribution, which serves as the starting point for the denoising process. As the process progresses, the right side of the plot showcases the generated data distributions from 4 different 2D ICNs. The probability density distribution plotted after the reverse process also shows four distinct peaks corresponding to the different ICNs. Each peak represents the unique attributes of an individual ICN, reflecting the variability and reconstruction of the underlying structures within the data.

### 3.4 Quantitative analysis of subject-specific ICN generation

The correlations between ICA-generated and DDPM-generated ICNs offer valuable insights into how each model captures variability within and between subjects. To explore this, a correlation matrix is calculated for two ICNs across four subjects, examining two key cases. The first case focuses on cross-correlation within the same subject, examining ICNs generated from different subsets of time points in the rs-fMRI data. The data for each subject, consisting of 490 time points, is divided into two equal parts: the first 245 and the last 245 time points. By comparing ICNs extracted from these subsets, we calculate a single cross-correlation value for each subject. This analysis reveals the consistency of ICNs within a subject across temporal segments, demonstrating their ability to capture within-subject variability in ICN patterns. For ICN 3, the within-subject correlations at different time points were high, with average values of 0.84 for Subject 1, 0.77 for Subject 2, and 0.85 for both Subjects 3 and 4, indicating that the DDPM effectively captures stable subject-specific features over time. This demonstrates the model’s ability to maintain consistency in the ICN structure for each subject over time. Meanwhile, for ICN 15, the within-subject correlations were 0.87, 0.90, 0.91, and 0.92, respectively, for each of the four subjects. These results demonstrate that ICNs derived from different temporal segments remain highly correlated, indicating that the variability captured by the models is not random but reflects meaningful patterns inherent to each subject.

Next, we calculate the cross-correlation between different subjects for the same network. This provides insights into the models’ capability to capture inter-subject variability. ICA directly computes ICNs for each of the four subjects, while DDPM averages results from 15 runs per subject to produce representative ICNs. A correlation matrix is then constructed, with entries indicating correlations between ICNs of different subjects as shown in Fig.5. Diagonal entries, which represent self-correlations, are excluded from the analysis. The average of the remaining correlations for each subject quantifies the similarity of that subject’s ICN to others. This approach allows us to evaluate the inter-subject consistency of ICNs, highlighting the models’ effectiveness in generalizing across individuals.

**FIGURE 5.**
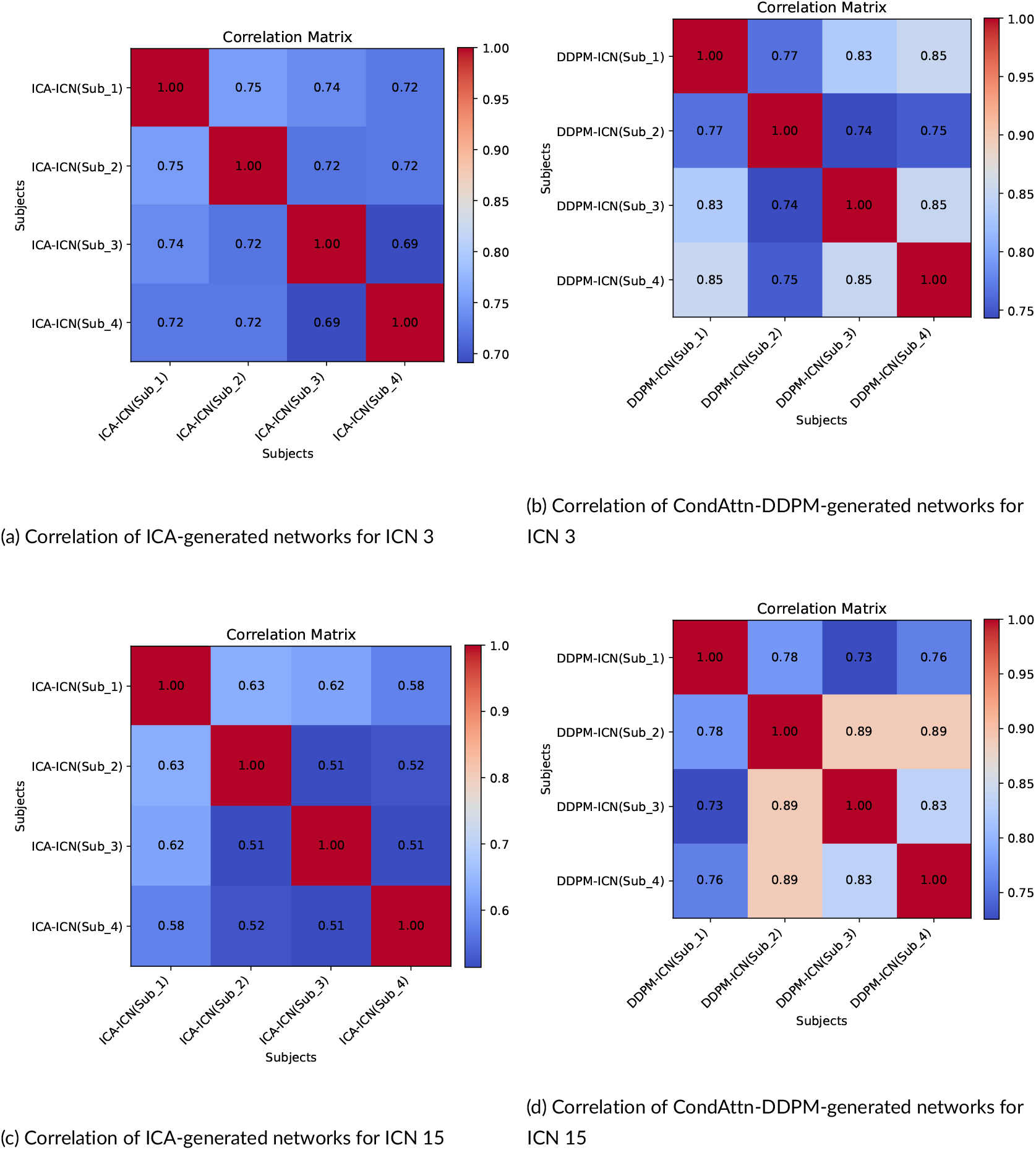
Comparison of ICA and CondAttn-DDPM frameworks in capturing between-subject variability using correlation matrices for ICN 3 and ICN 15. The analysis reveals that the CondAttn-DDPM framework consistently outperforms ICA in maintaining higher average correlations across subjects, indicating better preservation of subject-specific features. Specifically, for ICN 3, the average correlations for ICA were 0.74, 0.73, 0.72, and 0.71 for Subjects 1, 2, 3, and 4, respectively, while the CondAttn-DDPM framework achieved higher averages of 0.82, 0.75, 0.81, and 0.82 for the same subjects. Similarly, for ICN 15, the CondAttn-DDPM framework demonstrated superior performance with average correlations of 0.75, 0.85, 0.82, and 0.83 for Subjects 1, 2, 3, and 4, compared to ICA’s lower averages of 0.61, 0.56, 0.55, and 0.54 for the respective subjects. This highlights the enhanced ability of the CondAttn-DDPM framework to capture stable and meaningful patterns across different ICNs.

While comparing the correlations, the higher correlations within subjects compared to between subjects indicate that the model prioritizes subject-specific features while still capturing shared patterns of the ICN across individuals. This is important in neuroscience applications, as it demonstrates the model’s capacity to generate ICNs that retain individual uniqueness without losing the broader functional network structure.

Furthermore, Fig. 6 illustrates the differences in ICNs for a specific subject when generated using ICA and DDPM. Consistent with the correlation analysis, the visual comparison highlights that DDPM captures more individual-specific variability, while ICA emphasizes population-level consistency. Additionally, Fig. 7 presents a comparison across subjects, helping assess whether the methods capture consistent patterns or introduce subject-specific variability. In this comparison, DDPM showed greater variability across subjects, producing ICNs that reflect individual-specific characteristics, whereas ICA-generated ICNs showed more similarity across subjects, with only subtle differences.

**FIGURE 6.**
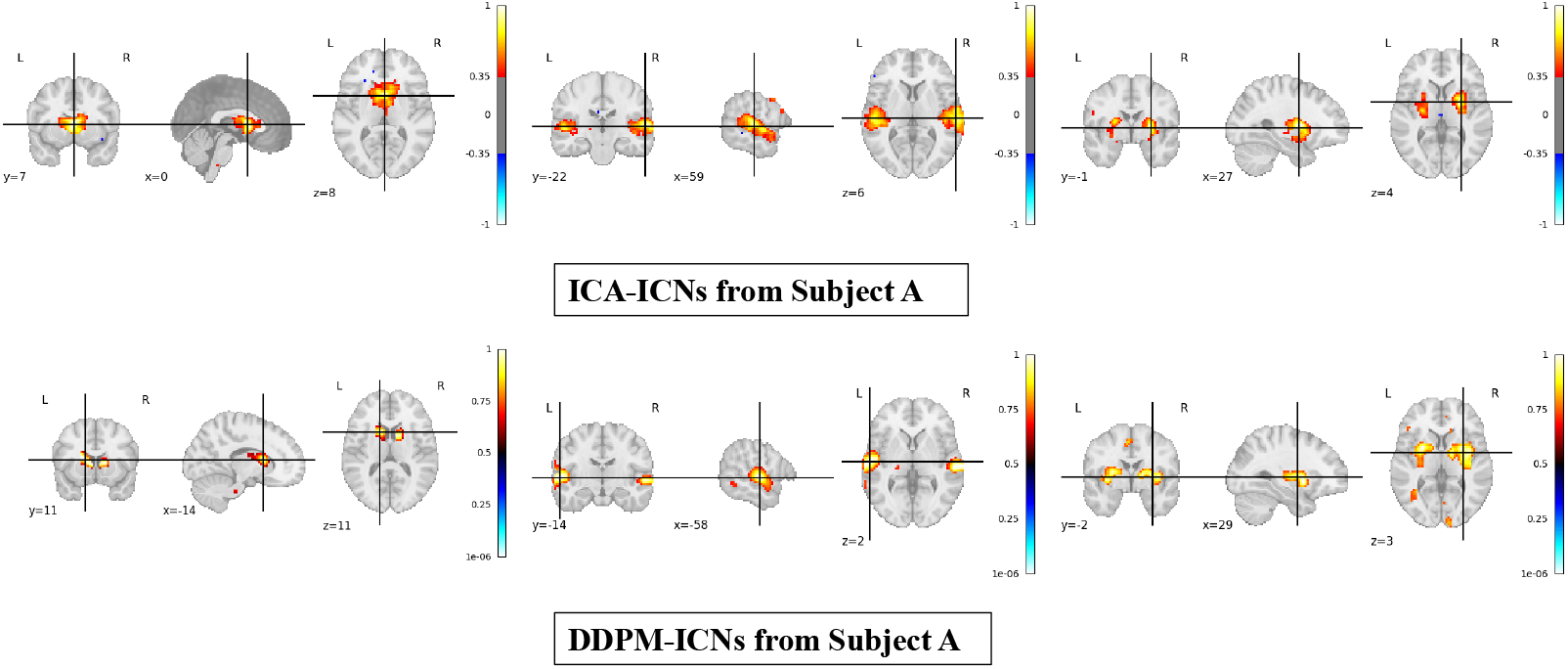
Comparison of ICNs generated using ICA and DDPMs for the same subject. ICA yields more stable and generalized ICNs, while DDPM produces ICNs that better reflect the subject’s unique connectivity pattern.

**FIGURE 7.**
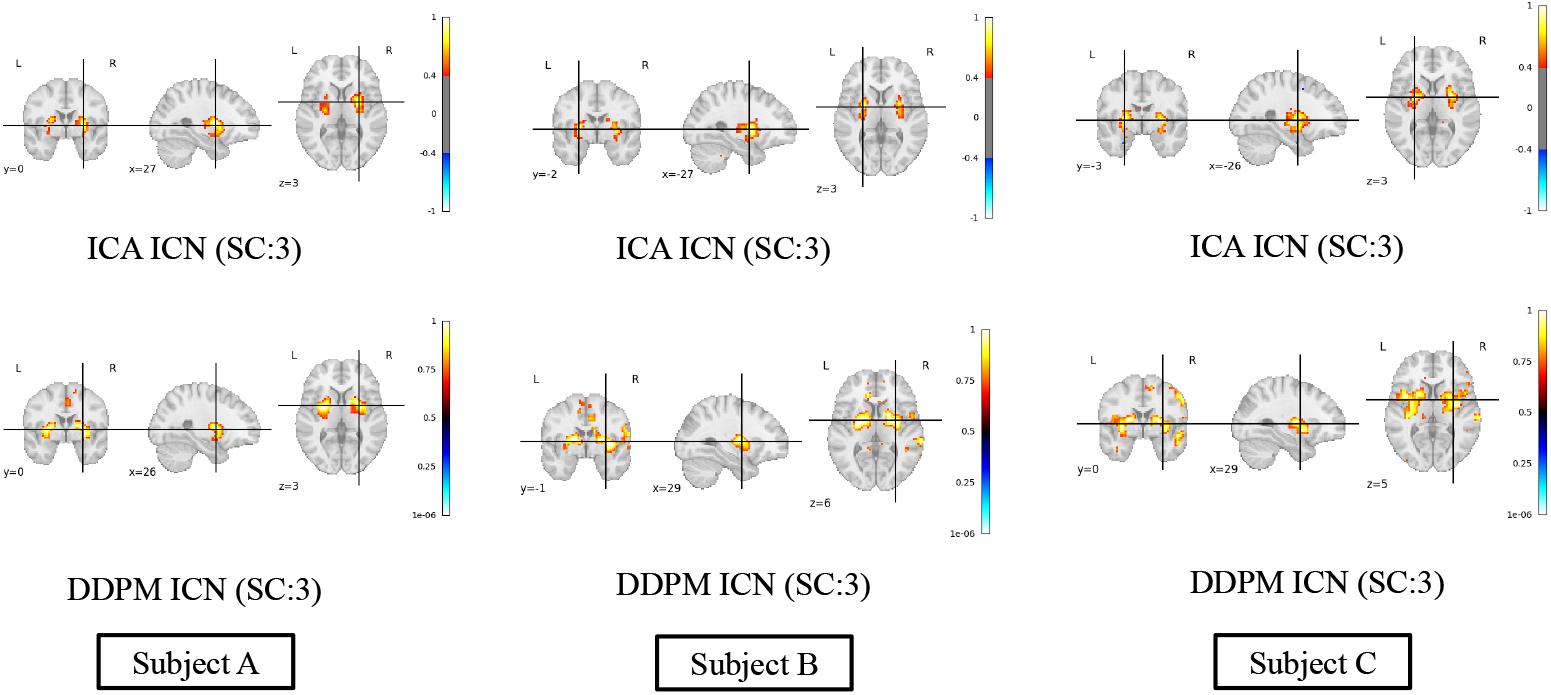
Comparison of ICNs generated for multiple subjects using ICA and DDPM. ICA-generated ICNs show less variation across subjects, indicating a focus on global, population-level features. DDPM-generated ICNs show more individual variation, capturing finer-grained connectivity differences unique to each subject.

## 4 DISCUSSION

This study presents two approaches for generating random and subject-specific ICNs using DDPM-based frameworks. To our knowledge, this is the first approach using diffusion models for blind source separation of fMRI data. ICA, although widely used in neuroimaging, is limited by its dependence on linear decomposition assumptions about independence and non-Gaussianity. This restricts its ability to extract the complex, non-linear relationships present in brain networks. Prior research has highlighted similar challenges with different types of ICA, especially for extracting intricate brain activity patterns [35, 36]. On the contrary, the proposed DDPM frameworks effectively capture non-linear relationships and generate ICNs with their adherent variability.

The experimental results also demonstrate the effectiveness of the proposed DDPM model in generating highquality, subject-specific ICNs, outperforming generative models like Progressive GANs in both visual quality and computational efficiency. While progressive GANs are known for high-resolution generation [37, 38], their artifacts, mode collapse, and limited variability hinder their application in complex neuroimaging data. In contrast, the iterative denoising mechanism in DDPMs enables the model to overcome these challenges. This allows for the generation of diverse ICNs without artifacts or random noise, thereby demonstrating the model’s robustness.

Furthermore, the model’s ability to maintain higher within-subject correlations than between-subject correlations demonstrates that the generated ICNs capture subject-specific variability, an essential component for personalized neuroimaging. When comparing the performance of the DDPM on ICN 3 and ICN 15, both networks exhibit strong within-subject stability across different time points. However, ICN 15 demonstrates slightly higher correlations (0.87–0.92) compared to ICN 3 (0.77–0.85), indicating that ICN 15 is better captured by the model for individual subjects. In terms of between-subject consistency, ICN 15 shows moderate-to-high cross-subject correlations (0.75–0.85), comparable to those observed for ICN 3 (0.75–0.82). Notably, ICN 15 displays a slightly wider range of cross-subject correlations, which may reflect greater subject-specific variability in its representation. Overall, these findings highlight distinct differences in the model’s ability to represent subject-specific variability and shared network features across ICNs. This result aligns with the growing emphasis on individualized brain network analysis for applications such as precision medicine, where understanding the unique neural configurations of patients can lead to tailored interventions [39, 40]. Recent studies have shown that personalized neuroimaging techniques improve diagnostic accuracy and treatment efficacy in various neurological and psychiatric disorders, reinforcing the necessity of capturing individual brain connectivity patterns [41]. Moreover, the reverse diffusion process, visualized through intensity distributions and histograms, showed the model’s ability to transform Gaussian noise into meaningful brain networks over time, reflecting the statistical properties of the original data. These results suggest that DDPMs not only capture variability across brain regions but also provide a promising framework for future studies focusing on subject-specific brain network modeling and clinical applications.

## 5 CONCLUSION AND FUTURE WORK

In conclusion, our work presents a novel framework that leverages the generative and nonlinear nature of DDPMs to synthesize high-quality 3D ICNs by training on a large and diverse dataset. These models address key limitations of traditional linear methods like ICA, specifically, their inability to capture non-linear relationships in brain activity. By integrating attention mechanisms into conditional DDPMs, the model enables the generation of subject-specific 3D ICNs. This innovative approach captures both within-subject and between-subject variability, providing more personalized insights into brain connectivity patterns. Unlike previous models restricted to 2D representations, our framework offers comprehensive 3D visualizations of ICNs. The model’s robustness and generalizability are validated through an external dataset, while comparative evaluations highlight its superiority over Progressive GAN models. Hence, providing individualized brain connectivity profiles using these models could aid in providing more precise personalized diagnostics and treatment planning. However, despite the promising advancements, our model has certain limitations. One key challenge lies in the computational complexity of generating subject-specific ICNs using CondAttn-DDPMs, which demand significant processing power and memory. This could limit the model’s scalability, particularly when working with larger datasets and higher-resolution images.

Additionally, while the integration of attention mechanisms enhances subject-specific ICN generation, the model’s performance may still be influenced by the quality and variability of the input data, potentially leading to inconsistencies in the synthesized ICNs. Our future work aims to address these limitations through several promising directions. First is the application of multimodal blind source separation techniques, integrating both fMRI and structural MRI (sMRI) data to provide richer representations of brain connectivity and improve the model’s robustness across varying conditions. Second, we aim to replace the U-Net backbone with a transformer-based architecture to improve the model’s ability to capture long-range dependencies in brain activity. Finally, to tackle the computational complexity of the DDPMs, future work will focus on learning to generate data in a compressed latent space and enabling fewer sampling steps than DDPMs.

## ACKNOWLEDGEMENTS

This work was supported by the Georgia State University RISE program, NSF grant 2112455, and NIH grants RO1MH123610 and RO1AG073949.

## DATA AVAILABILITY STATEMENT

All data of the UK Biobank database is subjected to third-party restrictions.

